# Consistency of SVDQuartets and Maximum Likelihood for Coalescent-based Species Tree Estimation

**DOI:** 10.1101/523050

**Authors:** Matthew Wascher, Laura Kubatko

## Abstract

Numerous methods for inferring species-level phylogenies under the coalescent model have been proposed within the last 20 years, and debates continue about the relative strengths and weaknesses of these methods. One desirable property of a phylogenetic estimator is that of statistical consistency, which means intuitively that as more data are collected, the probability that the estimated tree has the same topology as the true tree goes to 1. To date, consistency results for species tree inference under the multispecies coalescent have been derived only for summary statistics methods, such as ASTRAL and MP-EST. These methods have been found to be consistent given true gene trees, but may be inconsistent when gene trees are estimated from data for loci of finite length (Roch et al., 2019). Here we consider the question of statistical consistency for four taxa for SVDQuartets for general data types, as well as for the maximum likelihood (ML) method in the case in which the data are a collection of sites generated under the multispecies coalescent model such that the sites are conditionally independent given the species tree (we call these data Coalescent Independent Sites (CIS) data). We show that SVDQuartets is statistically consistent for all data types (i.e., for both CIS data and for multilocus data), and we derive its rate of convergence. We additionally show that ML is consistent for CIS data under the JC69 model, and discuss why a proof for the more general multilocus case is difficult. Finally, we compare the performance of maximum likelihood and SDVQuartets using simulation for both data types.

---

Advances in sequencing technology over the last 20 years have led to widespread availability of large-scale sequence data sets from multiple loci for which the goal is to obtain an estimate of the species-level phylogenetic relationships among the taxa under consideration. Analysis of such data has presented significant computational challenges, however, because inference methods must include models that capture variation at two distinct scales. First, a model for the process by which the phylogenetic histories of individual loci vary given the overall species tree must be developed. The coalescent process (Kingman, 1982b,c,a) is usually used for this purpose. Second, the mutation process arising along the locus-specific phylogenies, typically called gene trees, must be modeled. This is usually accomplished using standard nucleotide substitution models (Liò and Goldman, 1998). Together, these two model components are often referred to as the multispecies coalescent (MSC). Numerous methods for inference of species trees under the MSC have been developed (reviews of these methods can be found in several places, e.g., Liu et al. (2009) and Kubatko (2019)).

Inference of the species phylogeny under the MSC is challenging because the gene trees are not directly observed, and must therefore be integrated over when computing probabilities associated with the DNA sequence data. Consider a species tree with *M* species labeled 1, 2, …, *M*, and suppose that *m*_*j*_ individuals are sampled within each species *j*. Thus, 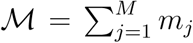 is the total number of sequences in the data set. Using the framework of the MSC, we denote the probability density of gene tree history *h* and associated vector of coalescent times **t**_*h*_, conditional on species tree topology *S* and vector of speciation times *τ*, by 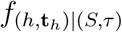 (see Rannala and Yang (2003) for a description of how to compute this density). We further define a site pattern to be an assignment of states 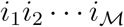 to the 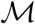 tips of the tree, such that *i*_*k*_ ∈ {*A, C, G, T*} for 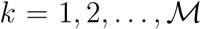, and we denote the probability of site pattern 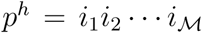 arising from *gene tree history* (*h*, **t**_*h*_) by 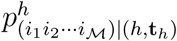. This probability is the usual phylogenetic likelihood along a gene tree, computed assuming one of the standard nucleotide substitution models. The probability of observing site pattern 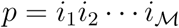 from the *species tree* is then given by

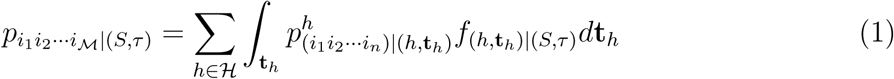

where the sum is taken over all gene tree histories *𝓗* with corresponding branch lengths **t**_*h*_ appropriately integrated out. See Chifman and Kubatko (2015) for full details of the calculations.

Note that Equation (1) implies that each site in the sequence alignment is an independent observation from the model; that is, each site represents a draw from the distribution of gene trees given the species tree as specified by the MSC, with subsequent mutation along the sampled gene tree according to one of the standard nucleotide substitution models. We use the term *coalescent independent sites* (CIS) to distinguish data of this type from SNP data, which do not usually include invariable sites. Under this model, a sample of *N* CIS can be viewed as a sample from the multinomial distribution, where the number of categories is the number of possible sites patterns, 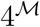, and the category probabilities are given by the site pattern probabilities. Thus, assuming that the sites are independent conditional on the species tree, the log likelihood of species tree (*S, τ*) is given by

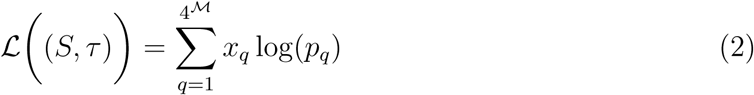

where *x*_*q*_ is the observed number of sites with pattern *q*, *p*_*q*_ is the probability of site pattern *q* under the model, 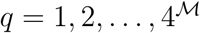, and 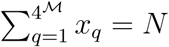. We note that the site pattern probabilities are functions of the parameters in the MSC model, including both the branch lengths and the effective population sizes along each branch. This likelihood has been mentioned earlier by Xu and Yang (2016).

The likelihood for multilocus data is more complicated, because in that case sites within a locus share the same gene tree and are thus correlated with one another, unless we condition on the gene tree. Suppose that there are *G* loci and that locus *g* has length *n*_*g*_. Let 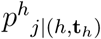 denote the probability that the site pattern observed for site *j* within a particular locus arises from gene tree history (*h*, **t**_*h*_). Then, the multilocus likelihood of species tree 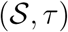 is

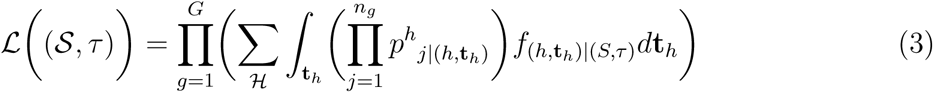

The outermost product is taken over the *G* loci, and assumes that the loci are independent, conditional on the species tree. Comparing the terms inside this outer product to the expression in Equation (1), we see that within the integral over the gene tree branch lengths, the product over the *n*_*g*_ sites within each gene must be taken. These sites are conditionally independent given the gene tree and branch lengths, but are not independent when conditioning only on the species tree because they share a common gene tree. The product appearing inside the integral makes it difficult to apply standard asymptotic arguments to this expression. Even taking the log of this likelihood, which allows the likelihood based on CIS data (Equation (2)) to be handled in a straightforward way, does not resolve the problem of the product appearing inside the integral. This also makes clear why computation of the species tree likelihood for multilocus data under the MSC model is challenging. In fact, we do not know of any direct implementations that compute this likelihood for trees larger than the four-taxon case we consider here.

To study the convergence properties of SVDQuartets and of maximum likelihood (ML), we consider the case in which *M* = 4 and *m*_*j*_ = 1 for all *j*, that is, we consider four-taxon trees with one sequence sampled in each species. In this case, there are 4^4^ = 256 possible site patterns, 15 rooted species trees, and 3 unrooted species trees. When considering ML to estimate the species tree, we restrict our attention to CIS data and use the likelihood given in Equation (2), given the difficulty in handling the multilocus likelihood discussed above. To find the ML estimate of the species tree for CIS data, one needs to be able to compute the true site pattern probabilities for each possible species tree. Formulas for these site pattern probabilities were given by Chifman and Kubatko (2015) for simple substitution models (e.g., JC69 (Jukes and Cantor, 1969)). Under the JC69 model and using these formulas with a single value of the effective population size parameter, *θ*, specified for the entire tree, we can find the ML estimate of the species tree by considering each of the 15 rooted species trees and finding the set of speciation times that maximize the likelihood for each. The tree with the highest likelihood is the ML estimate. We have implemented this method in R using the optim function to carry out the optimization for each topology. Our code can be found at https://github.com/lkubatko/SpeciesTreeConsistency.

To obtain an estimate of the four-taxon species tree for SVDQuartets for any data type (both CIS and multilocus data) and for the GTR+I+Γ model or any sub-model, let *L* denote the set of four taxa under consideration, and suppose that *L* is partitioned into two sets, *L*_1_ and *L*_2_, such that |*L*_1_| = |*L*_2_| = 2. We say that *L*_1_|*L*_2_ is a *split*. The split *L*_1_|*L*_2_ is *valid* for tree *S* if the subtrees containing the taxa in *L*_1_ and in *L*_2_ do not intersect; otherwise the split is not valid. For example, consider the tree ((1, 2), (3, 4)). The split 12|34 is valid, while the splits 13|24 and 14|23 are not valid.

For each of the three possible splits, the 256 possible site patterns can be arranged into a 16 × 16 matrix in which the rows of the matrix correspond to possible states for the taxa in *L*_1_ and the columns correspond to possible states for the taxa in *L*_2_. Such a matrix is called a *flattening matrix*, and is denoted 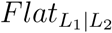. For an empirical data set, the entries of the matrix are the observed frequencies of the site pattern that corresponds to the row and column indices, i.e.,

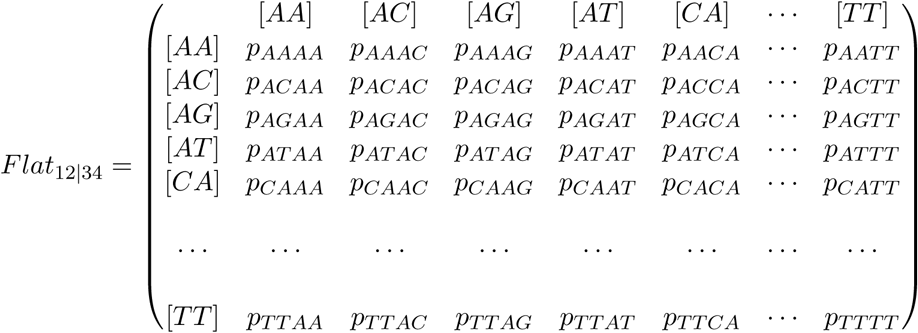

For example, the (3, 2) entry, *p*_*AGAC*_, is the probability of observing nucleotide *A* for taxon 1, *G* for taxon 2, *A* for taxon 3, and *C* for taxon 4. When the rows and columns of the matrix correspond to a valid split, the matrix will have rank 10 for data observed perfectly from the model. When the rows and columns correspond to a split that is not valid, the >matrix will be rank 16. The SVDQuartets method constructs three matrices (one for each of the three possible splits for four taxa), and computes the *SVD score* for each matrix,

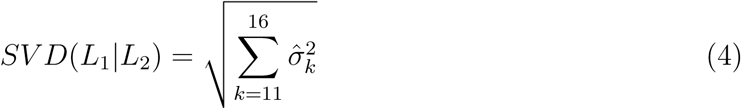

where 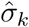 is the *k^th^* singular value computed for the matrix of observed site pattern frequencies. For observed data, the magnitudes of the 11^*th*^ through 16^*th*^ singular values are expected to be small when the matrix corresponds to the valid split, and thus the split *L*_1_|*L*_2_ with the lowest *SV D*(*L*_1_|*L*_2_) is selected. Note that in the case of four taxa, identifying the valid split is equivalent to inferring the unrooted species tree.

Under either criterion for estimation, we denote the estimator of the species tree by *S∗* and the true species tree by *S*. Intuitively, consistency means that as more data are used to form the species tree estimate, the probability that *S∗* = *S* goes to 1. SVDQuartets has been assumed to be statistically consistent, but a formal proof has not been provided. ML is known to be consistent when used to estimate gene trees, but consistency of ML has not been formally examined in the species tree case. In the sections below, we prove that SVDQuartets is consistent for both CIS and multilocus data and that ML is consistent for CIS data. We derive bounds for the error probability of SVDQuartets, and compare both methods using both theory and simulations.

## Consistency Results

We first define a generative model for multilocus data.

### Definition 0.1.

We assume that data are generated from the following statistical model:

1. Population and genome sizes are large enough that the fact that genes and sites are sampled without replacement can be ignored.
2. Define **p** to be a vector of multinomial probabilities such that if we select a gene at random and sample one nucleotide at random from that gene, the unconditional site pattern distribution **X** ∼ *Multinomial*(1, **p**).
3. Define 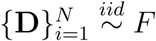 such that *E*(**D**_**i**_) = **0** and if we select *N* genes at random, each of the *N* genes will have multinomial site pattern probabilities **p**_**i**_ where 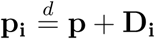.
4. Conditional on 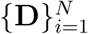, if **X**_**i**_ are the observed site pattern counts for a sample of *n*_*i*_ nucleotides from gene *i*, then **X**_**i**_ ∼ *Multinomial*(*n*_*i*_, **p**_**i**_) and the collection 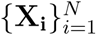 is independent.

Note that when *n*_*i*_ = 1 for all *i*, this model generates CIS data, which may thus be considered a special case of multilocus data.

### Consistency for Maximum Likelihood for CIS Data

While the literature contains numerous proofs of consistency of ML for estimation of gene trees (e.g., Yang (1994); Rogers (1997); RoyChoudhury et al. (2015); Truszkowski and Goldman (2016), described further below), no such proofs have been given for the case of ML estimation of the species tree, in part because it is not computationally feasible to use ML for species tree estimation under the coalescent model for trees of arbitrary size as discussed above. Some recent attention has also been given to evaluating the consistency of methods other than ML for estimating the species tree, but such work has focused primarily on the case in which multilocus data are collected and summary statistics methods are used to form estimators (Roch et al., 2019) or on the concatenation method (Roch and Steel, 2015). In this section, we formally prove that for CIS data ML estimation of the species tree under the multispecies coalescent described above is statistically consistent for four-taxon trees. We follow the proof of Truszkowski and Goldman (2016) for the case of gene trees, as most of their proof generalizes directly to the species tree case and their proof corrects the omissions of earlier proofs. We refer the reader to Truszkowski and Goldman (2016) for many of the details.

We first review related work for the case of ML estimation of gene trees, i.e., trees estimated using data from a single locus under one of the standard models of nucleotide substitution. Early proofs of the consistency of ML estimation for gene trees were given by Yang (1994) and Rogers (1997), but more recent examinations by RoyChoudhury et al. (2015) and Truszkowski and Goldman (2016) have found that these proofs are incomplete. RoyChoudhury et al. (2015) explains the problems with these proofs succinctly; we outline their argument here as it will apply to our proof for the species tree case given below. First, note that, assuming identifiability of the gene tree topology, which requires non-zero internal edges (for a proof, see, e.g., Allman et al. (2008), and note condition (2) in Section 2.1), the following proposition results from a straightforward application of the Strong Law of Large Numbers.

#### Proposition 0.2.

Suppose *T*_0_ is the true tree and *T*_*j*_ is any other tree. Then there exists *N* such that for all *n* ≥ *N*

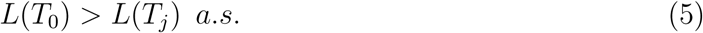

Though it is tempting to use Proposition (0.2) to claim consistency of the ML estimate of the gene tree topology, as noted by RoyChoudhury et al. (2015) and Truszkowski and Goldman (2016), this result is not sufficient to conclude that ML estimation is consistent. To see why, consider the typical definition of consistency of the maximum likelihood estimator (MLE) that states that if 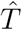 is the MLE then

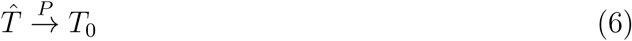

under a metric *D*(⋅, ⋅), where 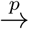 denotes convergence in probability. In order to guarantee that (6) holds, we either need to show that for any *ϵ* > 0 there exists some constant *C*_*ϵ*_ > 0 such that

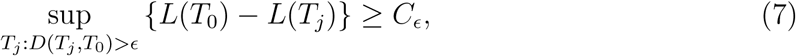

so that we are assured there cannot be trees of arbitrarily high likelihood far away from the true tree, or that the parameter space is compact. Under any reasonable metric, it is easy to see that the parameter space is not compact because it does not include trees with branches of length 0, as noted above. Truszkowski and Goldman (2016) provide a corrected proof by defining the following metric and showing that (7) holds for this metric (see Lemma 3 of Truszkowski and Goldman (2016)).

#### Definition 0.3

(Distance between two trees, Truszkowski and Goldman (2016)). For two taxa *a* and *b* in tree *S*, define their distance, *d*_*S*_(*a, b*), to be the sum of the lengths of all edges on the path from *a* to *b*. Further, define the distance between two trees *S*_1_ and *S*_2_ to be 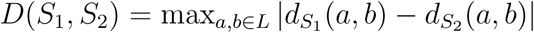. Note that *D*(⋅, ⋅) is a metric as long as all branch lengths are positive.

We now state and prove a modified version of Truszkowski and Goldman (2016)’s gene tree consistency result for the case of species trees estimated from a sample of CIS obtained under the multispecies coalescent.

#### Theorem 0.4

(Consistency of the ML estimator of the species tree for CIS data). *Let* 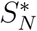 *denote the MLE of species tree S for a sample of N CIS obtained under the multispecies coalescent. Then* 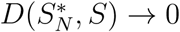 *with probability 1 as N → ∞*.

Our proof follows the general outline given by Truszkowski and Goldman (2016) for the gene tree case. Two crucial steps in their proof must be verified for the species tree case. First, the species tree must be identifiable, which has been established by Chifman and Kubatko (2015) for species trees that satisfy the molecular clock and by Long and Kubatko (2019) for non-clock species trees and for trees in which the effective population sizes vary throughout the tree. Second, a particular function of the pairwise distribution of states at the tips must satisfy a concavity condition. We state and verify this condition in the following proof.

*Proof*. Because the site pattern counts for a random sample of *N* CIS follow a multinomial distribution with probabilities given in Equation (1) above, the likelihood function for the ML estimate of the species tree is similar in form to that in the case of a gene tree. Thus Proposition (0.2) and most steps in the consistency proof given by Truszkowski and Goldman (2016) can be verified in a straightforward manner. The only non-trivial condition to be verified in the species tree case is that Lemma 3 of Truszkowski and Goldman (2016) still holds for the particular site pattern probabilities that arise in the species tree setting. This lemma involves some conditions on the pairwise site pattern probabilities, which we define below.

Following the notation of Truszkowski and Goldman (2016), let 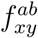 denote the frequency with which taxon *a* is observed to have state *x* and taxon *b* is observed to have state *y*, where *x, y* ∈ {*A, C, G, T*}. Let 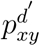 denote the probability that taxon *a* has state *x* and taxon *b* has state *y*, where *x, y* ∈ {*A, C, G, T*}, when *d*_*S*_(*a, b*) = *d′*. To verify Lemma 3 of Truszkowski and Goldman (2016) it is sufficient to verify that the function

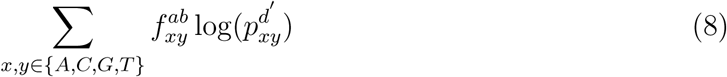

is concave in *d′*. Under the JC69 model (Jukes and Cantor, 1969) and the multspecies coalescent model, Chifman and Kubatko (2015) (see their Supplement A) gave explicit formulas for the site patterns probabilities on four-taxon trees. Using these with *µ* = 4/3 as specified by the JC69 model and *θ*, the effective population size parameter, set to 0.01, we can sum over pairs of taxa to find

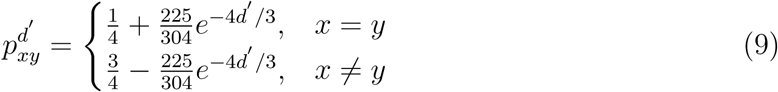

Ruskino (2018, personal communication) has derived more general expressions as a function the *θ* parameter that follow the same general form. Using these expressions, it is straightforward to verify that the expression in Equation (8) is concave in *d′* and thus that Lemma 3 of Truszkowski and Goldman (2016) holds. This establishes Theorem 0.4, and thus the ML estimate of the species tree for four taxa is statistically consistent for CIS data.

We next consider consistency for SVDQuartets.

### Consistency and Error Rate for SVDQuartets

Recall that for the SVDQuartets method, we choose the tree with split argmin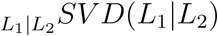 as our estimate of the species tree. In this section, we prove that this estimator is consistent for multilocus data in the following sense and give its rate of convergence.

#### Theorem 0.5

(Consistency of SVDQuartets). *Suppose that the conditions of the model proposed by Chifman and Kubatko (2015) are satisfied, and L*_1_|*L∗*_2_ *is the true valid split among splits with |L*_1_| = |*L*_2_| = *L*_2_. *Fix ϵ* > 0. *Assume* lim_*N→∞*_ max_*i*=1…*N*_ {*n*_*i*_} = *K* < ∞, *and that all of the entries of the vector* **p** *are strictly between* 0 *and* 1. *Then ∃N*_*c*_ *such that* 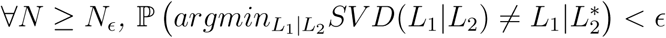.

We give the details of the proof of Theorem 0.5 in the remainder of this section. The result follows from consistency of the 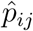 in the flattening matrix and the fact that singular values of a matrix satisfy a Lipschitz condition with respect to perturbations of the matrix (see Golub and VanLoan (2013)). The assumption that all of the entries of the vector **p** are strictly between 0 and 1 may seem arbitrary, but if this is not true, then we are considering the problem of estimating site pattern probabilities for sites that either always or never occur, and such cases are neither realistic nor interesting.

#### Lemma 0.6.

*[Corollary 8.6.2 of Golub and VanLoan (2013)] Let A, E ∈* ℝ^*m×n*^ *with m* ≥ *n*, *and let σ*_*i*_, *i* ∈ {1, … *n*}, *denote the singular values in descending order*. *Then for i* ∈ {1, … *n*}, |*σ*_*i*_(*A* + *E*) − *σ*_*i*_(*A*)| ≤ ||*E*||_2_ = *σ*_1_(*E*).

We first establish that the *p*_*ij*_> are consistently estimated (for two reasonable uses of multilocus data) and give their asymptotic error. The estimator 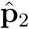 is currently used by SVDQuartets as implemented in PAUP* (Swofford, 2019).

#### Lemma 0.7.

*Suppose data are generated as in Definition 0.1 with N and n*_*i*_, *i* = 1 … *N*, *such that* lim_*N→∞*_ max_*i*=1…*N*_ {*n*_*i*_} = *K* < ∞, *and that all of the entries of the vector* **p** *are strictly between* 0 *and* 1. *Consider the estimators*

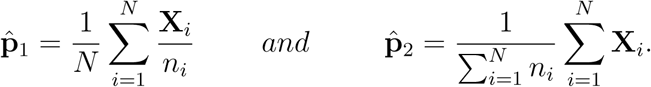

*The following hold*:

1. *Let* 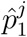 be the *j*^*th*^ *entry of* 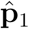, *and let p*^*j*^ *be the j*^*th*^ *entry of* **p** *in Definition 0.1. Then for any ϵ > 0*,

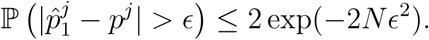
2. *Let* 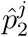 *be the j*^*th*^ *entry of* 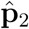. *Then for the K defined in Theorem 0.5 and ϵ > 0*,

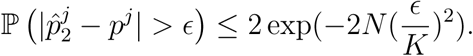

*Proof.* In both cases we will apply Hoeffding’s inequality.

1. Let *X*_*i,j*_ denote the *j*^*th*^ entry of **X**_*i*_, and let 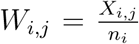. Then the *W*_*i,j*_ are bounded between 0 and 1 and independent with respect to the *i* index for any given *j*. Thus we can apply Hoeffding’s inequality to conclude

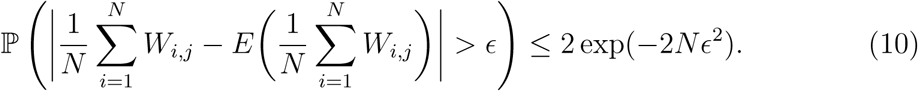 The first term in the expression above is 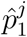. The second term is equal to 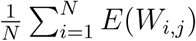. Now let *D*_*i,j*_ be the *j*^*th*^ entry of **D**_*i*_. Then

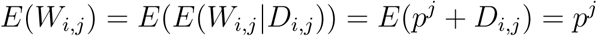

since our generative model assumes that *E*(**D**_*i*_) = **0**. Thus, 10 states that 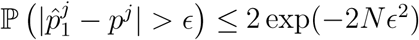 as desired.
2. Again let *X*_*i,j*_ denote the *j*^*th*^ entry of **X**_*i*_, and now let 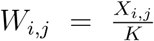 where *K* = lim_*N→∞*_ max_*i*=1*…N*_ {*n*_*i*_} which we have assumed is finite as stated in Theorem 0.5. Then the *W*_*i,j*_ are bounded between 0 and 1 and independent with respect to the *i* index for any *j*. Note that

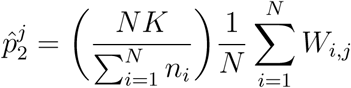

and

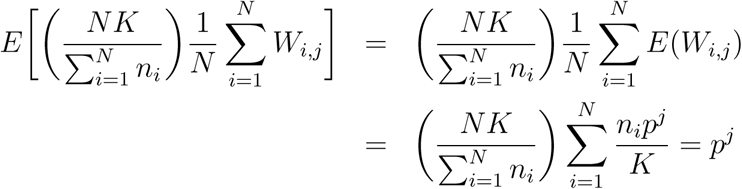

where the second equality above holds because 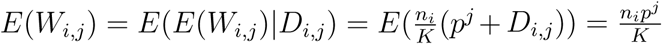. Thus, noting that 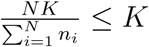, we have

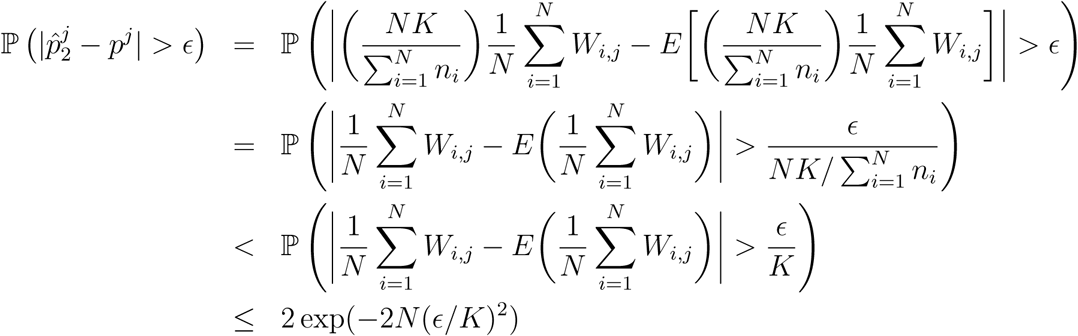

where the last inequality is the result of applying Hoeffding’s inequality. It might appear from the above bounds that 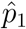 should be preferred to 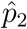 because the *K* term does not appear in the bound for 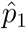, resulting in a smaller bound in that case. However, the bound in part (2) of the lemma is not tight. Rather, allowing the *K* term to appear in the exponent is simply a convenient way of dealing with the heterogeneity arising from the term 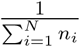. We discuss the relative merits of 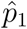 and 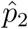 in the later section “SVDQuartets for Multilocus Data.”

If lim_*N→∞*_ max_*i*=1*…N*_ {*n*_*i*_} = *K* < ∞ does not hold, 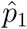 and 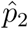 are still consistent estimators of **p**, but the deviations may have thinner tails than the bounds given above. Since it seems unrealistic that this assumption would be violated as in practice genes are finite in length, we do not provide a proof for the case where it does not hold.

#### Lemma 0.8.

*For any split L*_1_|*L*_2_, 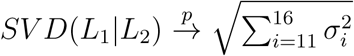 *where σ*_*i*_ *are the descending ordered singular values of the* 16 × 16 *matrix* 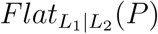.

*Proof.* Because *SV D*(*L*_1_|*L*_2_) is a continuous function of the vector (*σ*_1_, …, *σ*_16_), it suffices to show that 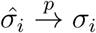 uniformly in *i*. Fix *ϵ, δ* > 0. We will show ∃*N*_*c,δ*_ such that ∀*N* ≥ *N*_*c,δ*_, 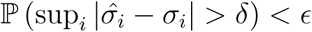.

We index the vector of site pattern probabilities **p** as {*p*_*ij*_} to match their locations in the flattening matrix. Note that this is a modification of the notation in Lemma 0.7 which used vectors to denote the site pattern probabilities. Likewise, we index 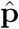 as 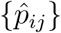. Define 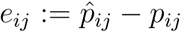, and observe that Lemma 0.7 implies that for any *i, j*, 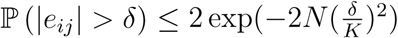. Now choose *N*_*ϵ,δ*_ large enough so that when 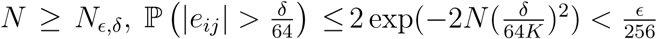. Using a union bound, we have

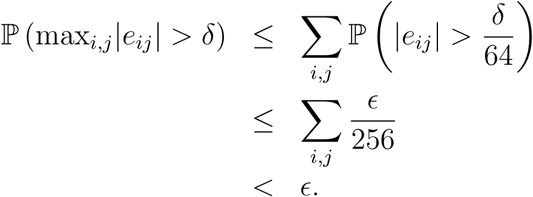

Now choose *E* in Lemma 0.6 to be *E* = *{e*_*ij*_}. Then 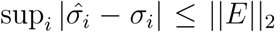. It is wellknown that for any matrix *E* ∈ ℝ^*k×k*^, 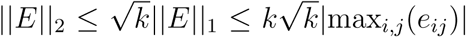. Applying this fact with *k* = 16, we have

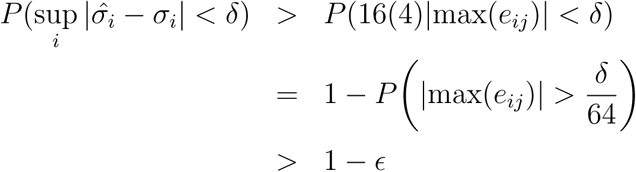

which gives 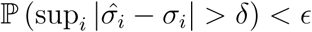, as desired.

We can now prove Theorem 0.5:

*Proof.* Theorem 1 of Chifman and Kubatko (2015) implies that 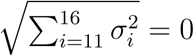 if and only if we choose the split *L*_1_|*L*∗_2_. Then because we have finitely many (3) splits to choose from, we can find some *c* > 0 such that 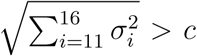 for any split *L*_1_|*L*_2_ ≠ *L*_1_|*L*∗. Fix *ϵ* > 0. Choose 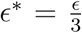 and 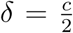. Then for the *N*_*c∗,δ*_ that satisfies Lemma 0.8 using *ϵ∗* and *δ*, for *N* ≥ *N*_*c∗,δ*_, we will have 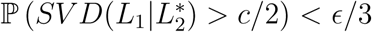 and 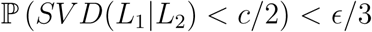 for every *L*_1_|*L*_2_ ≠ *L*_1_|*L∗*_2_. Then, using the union bound,

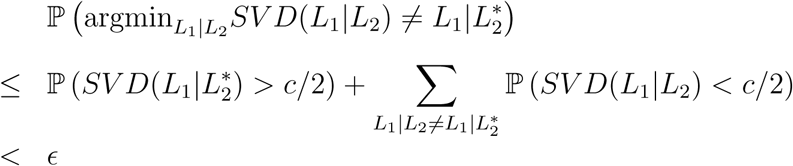

which completes the proof and establishes that SVDQuartets is a statistically consistent method for species tree estimation under the MSC.

We emphasize that this result proves consistency of SVDQuartets for both CIS data and for multilocus data using either of the estimators in Lemma 0.7 above. In both of these cases, the above result also gives a bound on the error rate, as described below.

#### Corollary 0.9.

*When estimating the split with SVDQuartets using a sample of n*_*i*_, *i* = 1 … *N*, *loci from each of N genes, there exists a constant σ*∗ > 0 *such that for large N the probability of choosing an incorrect split is bounded by*

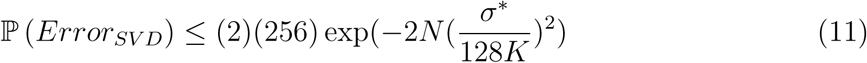

*Proof.* Let *σ*∗ be the smallest in absolute value of the 11^*th*^ −16^*th*^ nonzero singular values among all possible splits *L*_1_|*L*_2_. Note that as a consequence of Lemma 0.6, for any split and each 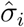,

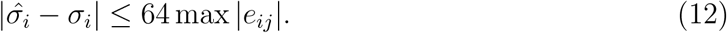

Let 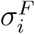 denote the *i*^*th*^ singular value for an incorrect split for any *i* = 11, …, 16, and let 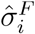 denote the corresponding observed value. Applying (12) and assuming that 64 max 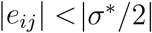 gives

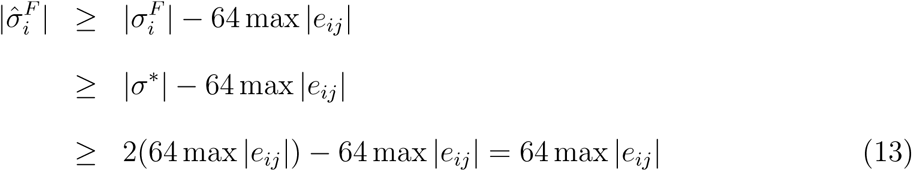

for each *i* = 11, …, 16.

Applying (12) again to singular values from the true split gives 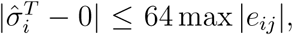 and we have

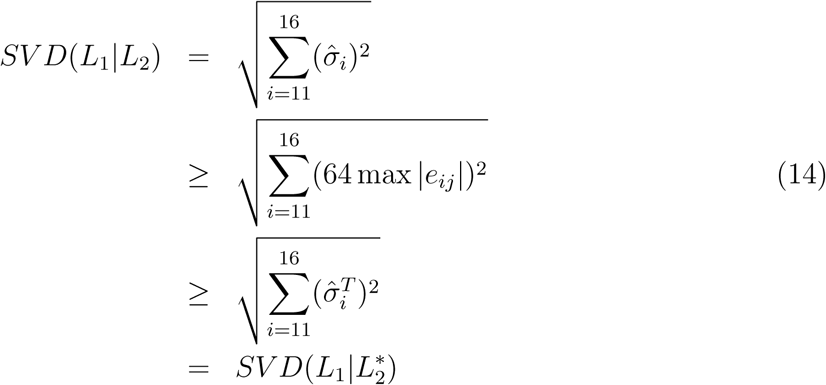

where (14) follows from (13). This establishes that the correct split will be selected by SVDQuartets whenever 64 max |*e*_*ij*_| < |*σ*∗/_2_|.

The probability that SVDQuartets makes an error in selecting the split can thus be given by

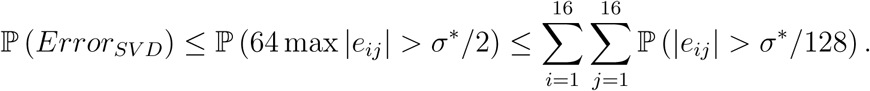

Recall from Lemmas 0.7 and 0.8 that 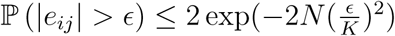, so

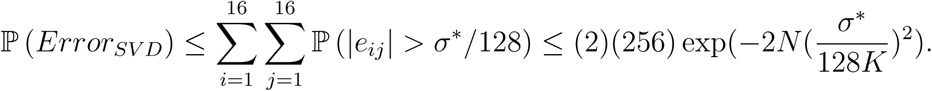

Note that our proof of consistency and error bound derivation depend on the structure of a four-taxon species tree only insofar as Theorem 1 of Chifman and Kubatko (2015) has only been proven for trees of four taxa and our choice of constants. Should that result be extended to trees with a larger number of taxa, our arguments above imply that the estimator based on the SVD score in such cases would also be consistent for multilocus data and would have an error rate bound of 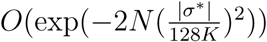.

### Comparison of Asymptotic Properties of ML and SVDQuartets

### Theoretical Comparison

Shi and Yang (2018) conjecture that SVDQuartets is inefficient compared to ML when both are applied to multilocus data, as measured by the probability of recovering the correct species tree. Note that this is a different notion of efficiency than that which is applied in typical statistical settings, raising the question of whether classical statistical results concerning asymptotic efficiency of ML estimators (see, e.g., Lehmann and Casella (1998)) apply in this case. As mentioned earlier in our discussion of consistency, it is not clear whether ML for the species tree estimation problem satisfies the general conditions of Wald (1949) for consistency. Nonetheless, we have been able to show that both ML for CIS data and SVDQuartets for multilocus and CIS data give statistically consistent estimators of the species tree. We next try to summarize what is know about the asymptotic error probabilities of the methods, as a way of addressing the claim made by Shi and Yang (2018) about the relative efficiency of the two methods.

To our knowledge, error rate bounds for ML when applied to multilocus data have been rigorously derived in only a few special cases. Xu and Yang (2016) showed that in the case of a three-taxon species tree, the probability of choosing the wrong topology when using ML for data consisting of rooted gene trees is approximately

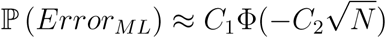

for explicit constants *C*_1_ and *C*_2_ that depend on the probabilities of the three possible gene trees that can arise within the species tree (which in turn can be computed from the other parameters.) It is important to note, however, that this result is an approximation rather than a bound. It does not account for the rate at which 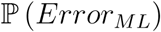, which is not exactly normal, converges to a normal distribution, and this rate could potentially be slower than the decay of the normal tail given by the approximating expression 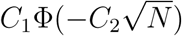. An equivalent result for four-taxon species trees has not been derived.

Another partial result about the error rate of ML estimation comes from the following idea. Suppose that rather than sample *n*_*i*_ sites from each of *N* loci, we are able to sample gene trees directly, so that we in fact know the topology and branch lengths of each of the *N* sampled gene trees, *G*_1_ … *G*_*N*_. Letting **p**_*l*_ denote the observed site patterns (i.e., the alignment) for gene *l*, we note that in this case,

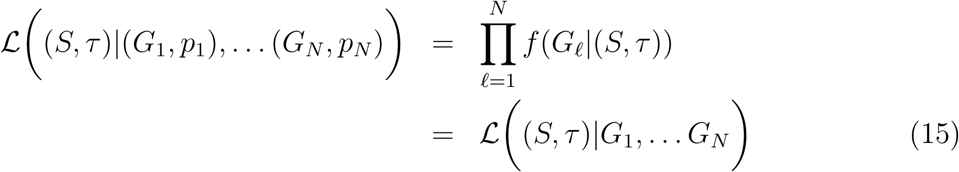

where *f* is the gene tree density under the MSC. In words, if we observe the gene trees directly, the alignments give no additional information and the likelihood of interest is the species tree likelihood based on the sampled gene trees. Furthermore, since this sampling scheme uses strictly more information than sampling only finite-length alignments, it seems reasonable to assume that its estimation power should be at least as high, (i.e., its error rate no worse than that of ML based on multilocus sampling) although this also requires a proof to be made fully rigorous.

Results about error rates for trees of any size have been derived for the problem of estimating the species tree topology using gene trees directly. Liu et al. (2010) showed that for their maximum tree method, the probability of choosing the wrong topology is bounded by an expression of the form

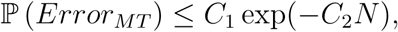

and that if all populations have the same size, the maximum tree estimator is also the ML estimator. This result is a rigorous upper bound rather than an approximation. We note that it is comparable to our result for SVDQuartets insofar as both bounds take the form *C*_1_ exp(−*C*_2_*N*), albeit likely for different values of *C*_1_ and *C*_2_.

A rigorous comparison of the performance of ML and SVDQuartets is inconclusive, in large part because not enough is known about the performance of ML. Since the result of Xu and Yang (2016) comes from multinomial probabilities, it is likely that applying Hoeffding’s inequality in that case would also yield a bound of the form *C*_1_ exp(−*C*_2_*N*) in addition to the approximation given in their work, although we have not rigorously verified this. One might additionally conjecture that such a result holds for species trees with arbitrary numbers of taxa, rather than just the three-taxon species tree. If this is true, then we could say that ML and SVDQuartets both have error rate bounds of the form *C*_1_ exp(−*C*_2_*N*), where the constants *C*_1_ and *C*_2_ likely differ between the methods, but we cannot compare beyond this statement. We hope that scholars interested in comparing the performance of ML and SVDQuartets will derive more complete rigorous results that will allow for a more comprehensive theoretical comparison.

### Comparison via Simulation

We conducted several simulation studies to comparatively evaluate the performance of ML and SVDQuartets. In the first simulation study, CIS data were simulated along the four-taxon symmetric and asymmetric species trees by first simulating gene trees using the package COAL (Degnan and Salter, 2005) and then simulating sequence data under the JC69 model (Jukes and Cantor, 1969) using Seq-Gen (Rambaut and Grassly, 1997). The JC69 model was used because Chifman and Kubatko (2015) provided explicit formulas for the site pattern probabilities for four-taxon trees under the coalescent for this model, allowing us to implement the maximum likelihood method in this case. We considered three species trees with all internal branch lengths and all external branch lengths leading to cherries set to the same value, either 0.5, 1.0, or 2.0 in coalescent units. We also consider three species trees with varying branch lengths. In these three cases, all branch lengths were either 0.5 or 1.0, and the placement of the shorter branches was varied between internal and external branches. The precise trees used are given in the captions to Figures 1 and 2, which show the simulation results. For all of the model trees, we set the effective population size parameter *θ* = 4*Nµ* to 0.001, 0.005, or 0.01 for all branches. In addition to examining the performance of SVDQuartets for CIS data, we examined its performance on SNP data by re-running it on each simulated dataset after removing all of the constant sites. Since SNP data are more commonly collected, this will provide an indication of how much information is lost in moving from CIS to SNP data when using SVDQuartets for inference.

**Figure 1:**
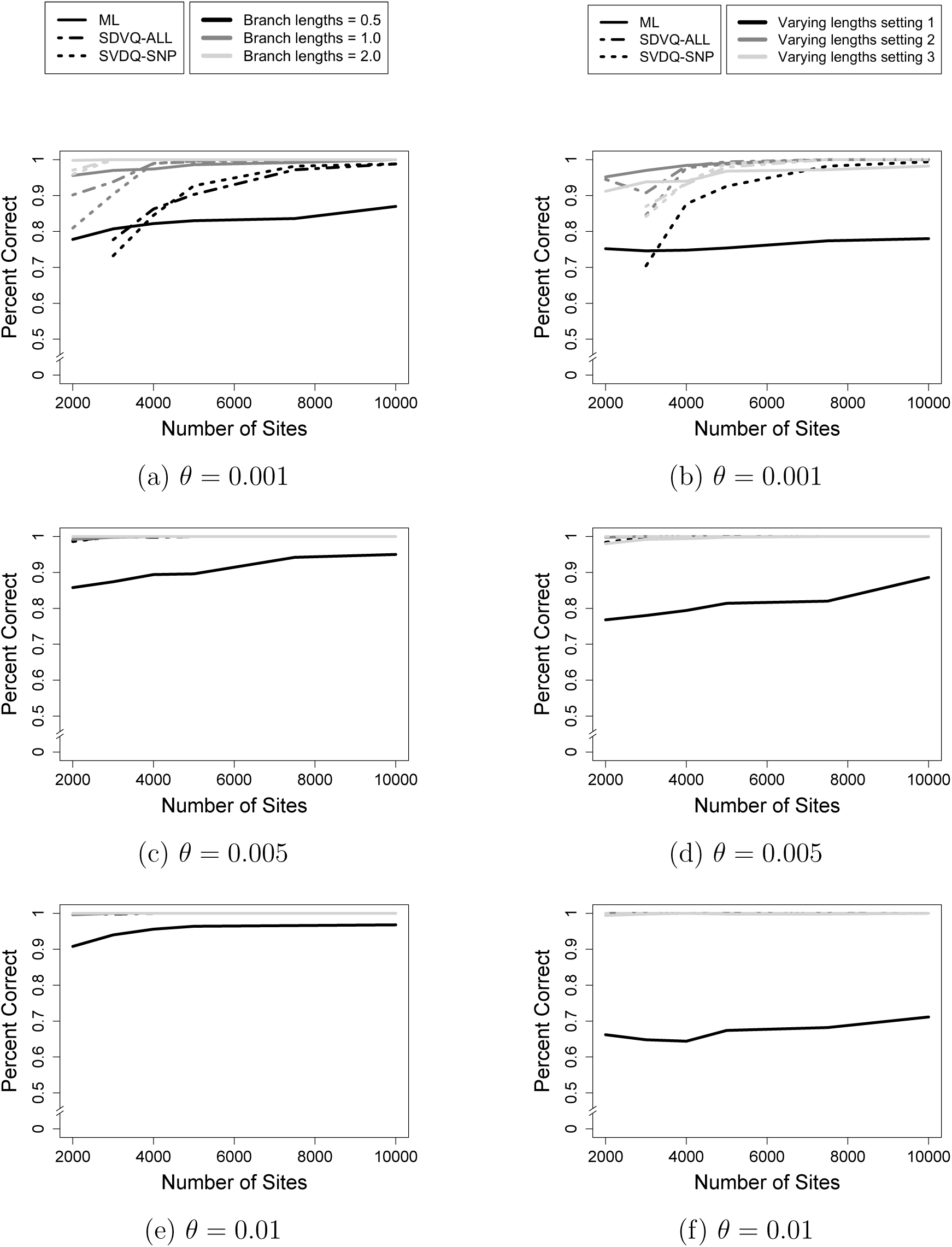
Results of the simulation study for the symmetric species tree for CIS data. The x-axis shows the number of CIS, and the y-axis shows the proportion of correctly-estimated unrooted species trees for each method. (a) *θ* = 0.001, all branch lengths equal to value given in the legend; (b) *θ* = 0.001, varying branch lengths; (c) *θ* = 0.005, all branch lengths equal to the value given in the legend; (d) *θ* = 0.005, varying branch lengths; (e) *θ* = 0.01, all branch lengths equal to the value given in the legend; (f) *θ* = 0.01, varying branch lengths. For (b), (d), and (f), setting 1 refers to tree ((Species1:1.0,Species2:1.0):0.5,(Species3:1.0,Species4:1.0):0.5); setting 2 refers to tree ((Species1:0.5,Species2:0.5):1.0,(Species3:0.5,Species4:0.5):1.0); and setting 3 refers to tree ((Species1:1.0,Species2:1.0):0.5,(Species3:0.5,Species4:0.5):1.0).

**Figure 2:**
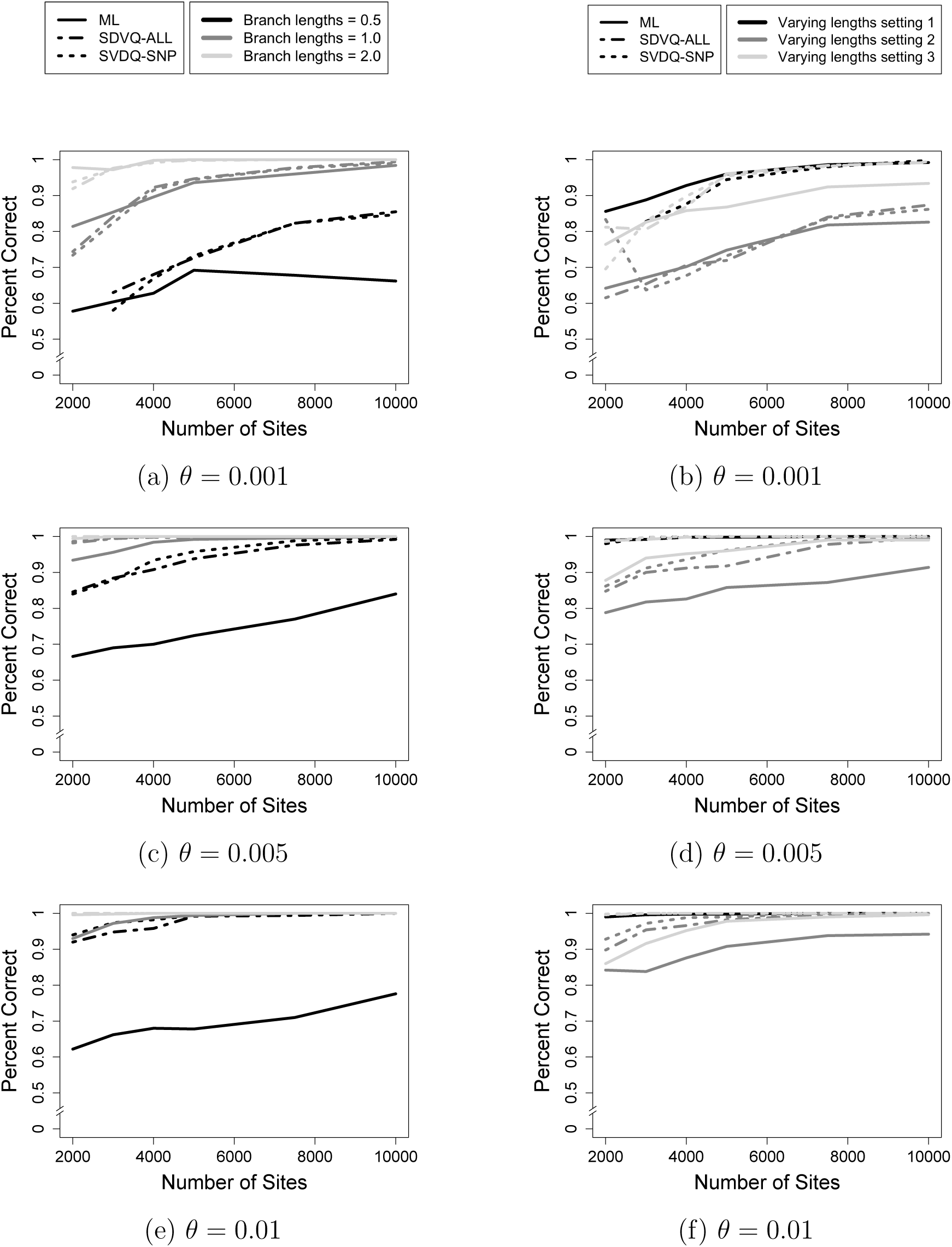
Results of the simulation study for the asymmetric species tree for CIS data. The x-axis shows the number of CIS, and the y-axis shows the proportion of correctly-estimated unrooted species trees for each method. (a) *θ* = 0.001, all branch lengths equal to value given in the legend; (b) *θ* = 0.001, varying branch lengths; (c) *θ* = 0.005, all branch lengths equal to the value given in the legend; (d) *θ* = 0.005, varying branch lengths; (e) *θ* = 0.01, all branch lengths equal to the value given in the legend; (f) *θ* = 0.01, varying branch lengths. For (b), (d), and (f), setting 1 refers to tree (Species4:2.5,(Species3:1.5,(Species2:0.5,Species1:0.5):1.0):1.0); setting 2 refers to tree (Species4:2.0,(Species3:1.0,(Species2:0.5,Species1:0.5):0.5):1.0); and setting 3 refers to tree (Species4:2.5,(Species3:2.0,(Species2:1.0,Species1:1.0):1.0):0.5).

For each of the three methods (SVDQuartets for CIS data, SVDQuartets for SNP data, and ML), we examined the proportion of times out of the 500 replicates that each of the methods correctly estimated the unrooted species tree when the total number of sites sampled ranged from 1,000 to 10,000. In some cases, particularly those in which the overall mutation rate is low, as often results from both small effective population size and short branches, the ML algorithm will not converge and/or singular values cannot be computed accurately enough to infer the tree with SVDQuartets. When this occurred, we discarded that replicate from the summary of that method’s performance. If a particular simulation setting had fewer than 100 replicates in which estimation was completed without error, we did not include the result for that setting in the relevant figure.

Our second simulation study considers multilocus data. We applied SVDQuartets as implemented in PAUP*, which ignores information about loci and treats the data as CIS data (this is the common and recommended practice for SVDQuartets at present). Because the multilocus likelihood is not computationally tractable, we approximated ML inference by running the BPP software (Yang and Rannala, 2014; Yang, 2015; Rannala and Yang, 2017; Flouris et al., 2018) with the prior for *τ* set to IG(3,0.015) and the prior for *θ* set to IG(3,0.01). We hereafter refer to these as the default priors. We discarded the first 400 samples as burnin, and recorded every other sampled tree for a total of 1,500 samples. For four-taxon trees, the species tree with the highest posterior probability will be generally equivalent to the ML tree. We consider the same model trees as in the first simulation study and the same choices of *θ*. We used 5, 10, 15, 20, 25, 35, or 50 loci, with 200bp per locus, and replicated each simulation condition 500 times. Replicates in which SVDQuartets failed to return an estimate of the species tree due to numerical imprecision of the singular value computation were discarded, as described above.

For the third simulation study, we considered more difficult species trees, namely those found in the anomaly zone. In particular, we considered the tree found in Xu and Yang (2016), which is given by (Species1:0.48,(Species2:0.44,(Species3:0.4,Species4:0.4):0.04):0.04) when branch lengths are reported in coalescent units. For the analysis in BPP, we considered both default priors, as we did in our second simulation study, and priors suggested by the simulation study in Xu and Yang (2016). Because the current version of BPP uses the inverse gamma (IG) distribution, rather than the gamma distribution used by Xu and Yang (2016), we use inverse gamma prior for *θ* and *τ* that have *α* = 3 and mean set to the true value. Another difference from our simulation conditions and the simulation carried out by Xu and Yang (2016) is the number of sites per locus. Thus, we include one set of simulations with 200bp per loci (as above) and another with 1000bp per loci as in Xu and Yang (2016). Our preliminary simulations with these settings indicated that many more sites were needed for accurate inference for both ML and SVQuartets, and thus for the CIS simulations, we considered the number of sites ranging from 10,000 to 1,000,000. For the multilocus simulations, we considered 50, 100, 150, 200, 300, and 400 loci. Because of the additional computational cost associated with the large number of sites, we used only 100 replicates of each simulation condition. Finally, we considered *θ* values that differed somewhat from those used above, in order to reproduce the results of Xu and Yang (2016) reported in their Figure 7. In particular, we considered *θ* = 0.05 (the value used by Xu and Yang (2016)) and *θ* = 0.01. As mentioned above, we considered both informative and default priors for BPP. For the informative priors, we assumed *τ* is IG(3.0,0.024), and *θ* is IG(3,0.1) when the true value of *θ* = 0.05 and *θ* is IG(3,0.02) when the true value of *θ* = 0.01.

Figure 1 shows the results of the first simulation for the symmetric species tree, and Figure 2 shows the results for the asymmetric species tree. In general, both methods are able to accurately infer the unrooted four-taxon species tree with sufficient data. When the model species tree is symmetric (Figure 1), both methods are very accurate when the branch lengths within the model species tree are equal, though shorter branch lengths and lower values of *θ* (settings which correspond to lower overall mutation rates) are more difficult for both methods. When branch lengths vary within the tree for the symmetric case, the first varying lengths setting, which corresponds to short internal branch lengths, was most difficult for both methods, though ML in general showed higher error than SVDQuartets for all three choices of *θ*. Results for the asymmetric model species tree (Figure 2) were likewise similar for both methods, with shorter internal branch lengths corresponding to lower accuracy for both methods. An important observation is that SVDQuartets does not decrease in accuracy when applied to SNP data as compared to CIS data. This can be explained by the observation that constant site patterns do not play a role in the reduced rank result of Chifman and Kubatko (2015) that is the basis for the SVDQuartets method. Figure 3 shows the results of the second simulation for the symmetric species tree, and Figure 4 shows the results for the asymmetric species tree. In most cases, the accuracy of SVDQuartets is lower than that of BPP, which is not surprising given that BPP is designed explicitly for multilocus data and SVDQuartets is designed for CIS data. It is clear that as the number of loci increases and the branches become longer, BPP accurately infers the true four-taxon species tree, while the performance of SVDQuartets lags behind. This suggests that the SDVDQuartets method may be most useful for genome-scale multilocus data, a setting in which the asymptotic consistency result suggests good performance and in which Bayesian methods become computationally expensive, while Bayesian methods such as BPP may be more appropriate when a more limited number of loci are available. We further compare the two frameworks (i.e., a likelihood-based framework such as BPP and the SVDQuartets methods) in the Discussion.

**Figure 3:**
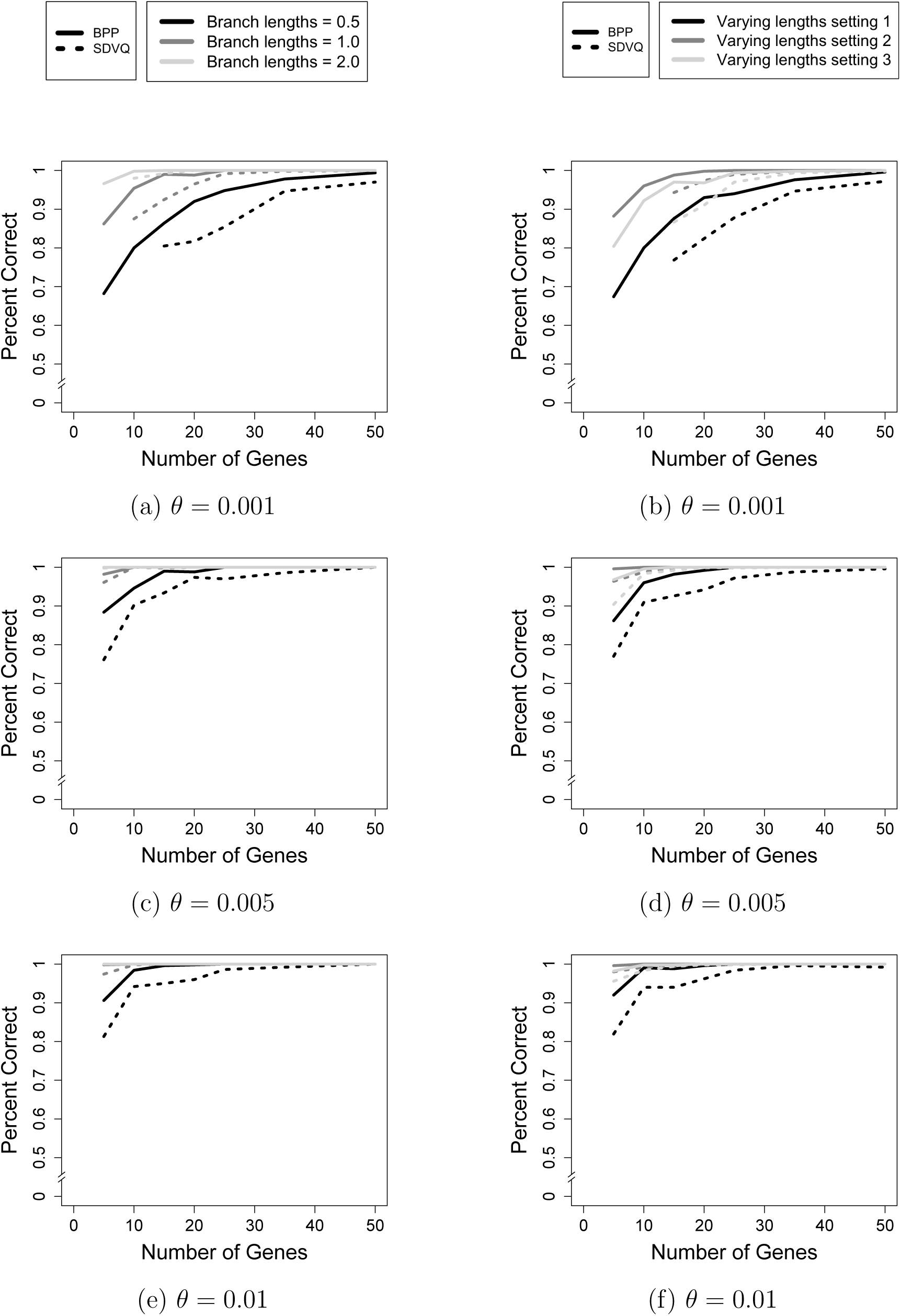
Results of the simulation study for the symmetric species tree for multilocus data. The x-axis shows the number of genes, and the y-axis shows the proportion of correctly-estimated unrooted species trees for each method. (a) *θ* = 0:001, all branch lengths equal to value given in the legend; (b) *θ* = 0:001, varying branch lengths; (c) *θ* = 0:005, all branch lengths equal to the value given in the legend; (d) *θ* = 0:005, varying branch lengths; (e) *θ* = 0:01, all branch lengths equal to the value given in the legend; (f) *θ* = 0:01, varying branch lengths. For (b), (d), and (f), setting 1 refers to tree ((Species1:1.0,Species2:1.0):0.5,(Species3:1.0,Species4:1.0):0.5); setting 2 refers to tree ((Species1:0.5,Species2:0.5):1.0,(Species3:0.5,Species4:0.5):1.0); and setting 3 refers to tree ((Species1:1.0,Species2:1.0):0.5,(Species3:0.5,Species4:0.5):1.0).

**Figure 4:**
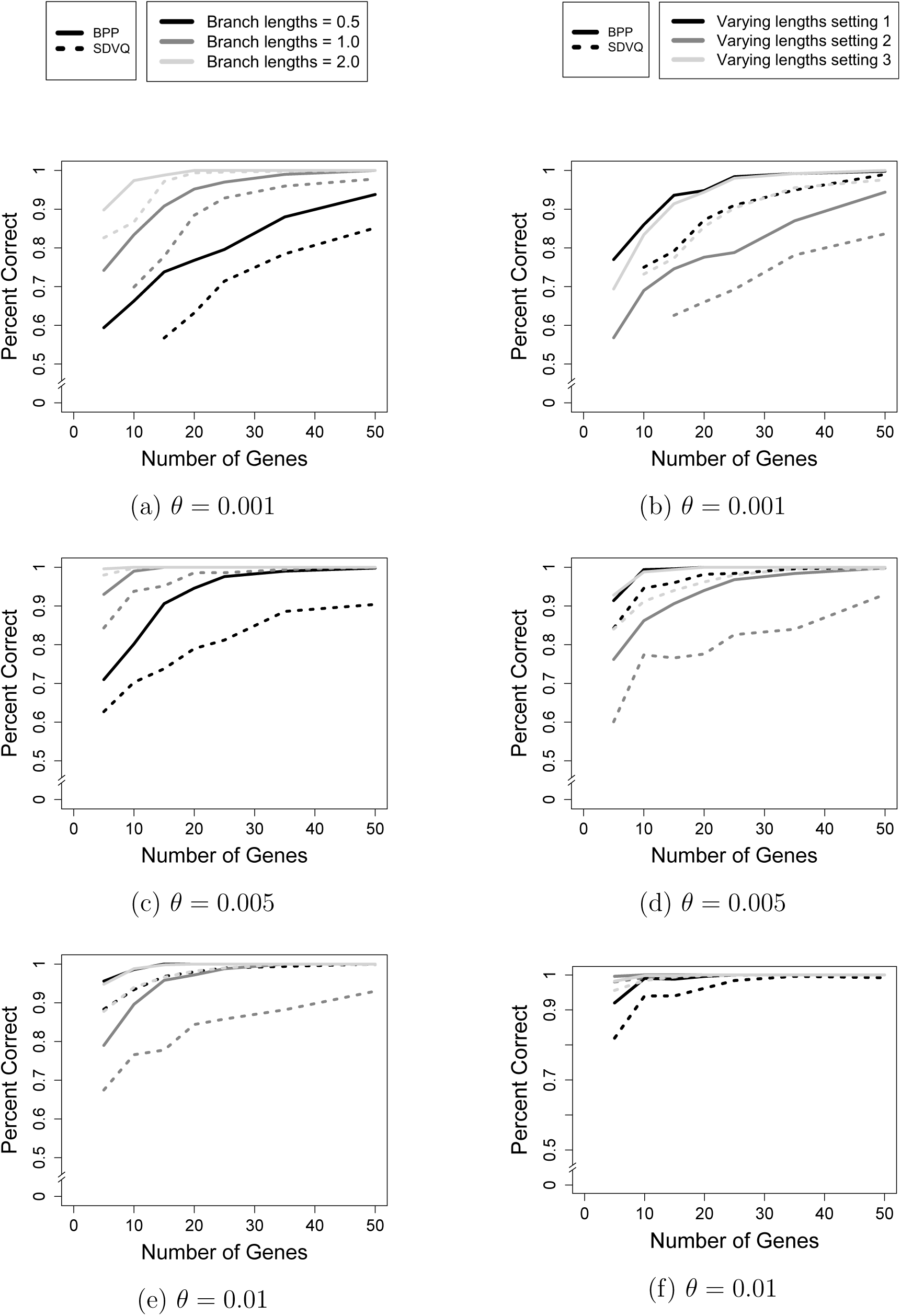
Results of the simulation study for the asymmetric species tree for multilocus data. The x-axis shows the number of genes, and the y-axis shows the proportion of correctly-estimated unrooted species trees for each method. (a) *θ* = 0:001, all branch lengths equal to value given in the legend; (b) *θ* = 0:001, varying branch lengths; (c) *θ* = 0:005, all branch lengths equal to the value given in the legend; (d) *θ* = 0:005, varying branch lengths; (e) *θ* = 0:01, all branch lengths equal to the value given in the legend; (f) *θ* = 0:01, varying branch lengths. For (b), (d), and (f), setting 1 refers to tree ((Species1:1.0,Species2:1.0):0.5,(Species3:1.0,Species4:1.0):0.5); setting 2 refers to tree ((Species1:0.5,Species2:0.5):1.0,(Species3:0.5,Species4:0.5):1.0); and setting 3 refers to tree ((Species1:1.0,Species2:1.0):0.5,(Species3:0.5,Species4:0.5):1.0).

The results of the third simulation study are shown in Figure 5 for both CIS data (first row) and multilocus data (second row). As expected, for species trees for which there are anomalous gene trees, estimation of the correct species tree is more difficult for both methods, and more data are required to achieve reasonable accuracy. In the case of CIS data, SVDQuartets performs well with accuracy near 100% once a sufficient amount of data are available, in this case more than 500,000 sites. It is again worth noting that the accuracy of the method applied to SNP data is nearly identical to the CIS case, suggesting that SVDQuartets will be a very effective method when genome-scale SNP data are available. The performance of ML lags behind, likely due to the rather low number of informative sites available when species tree branch lengths are extremely short. For example, when branch length are short and *θ* is not very large, most of the site patterns generated will be constant (i.e., invariable) site patterns. These site patterns are not informative for either SVDQuartets or ML. The next most frequently occurring site patterns will be those with a different nucleotide in only one species. Such site patterns provide no information about topology for ML, since they don’t provide information about which two taxa are most closely related. However, these site patterns are informative for SDVQuartets, because the reduced rank result on which the method is based uses the relationship that site patterns *xxxy* and *xxyx* should occur in equal frequency. Thus it is reasonable that SVDQuartets performs better than ML for CIS and SNP data in low-information settings such as this.

**Figure 5:**
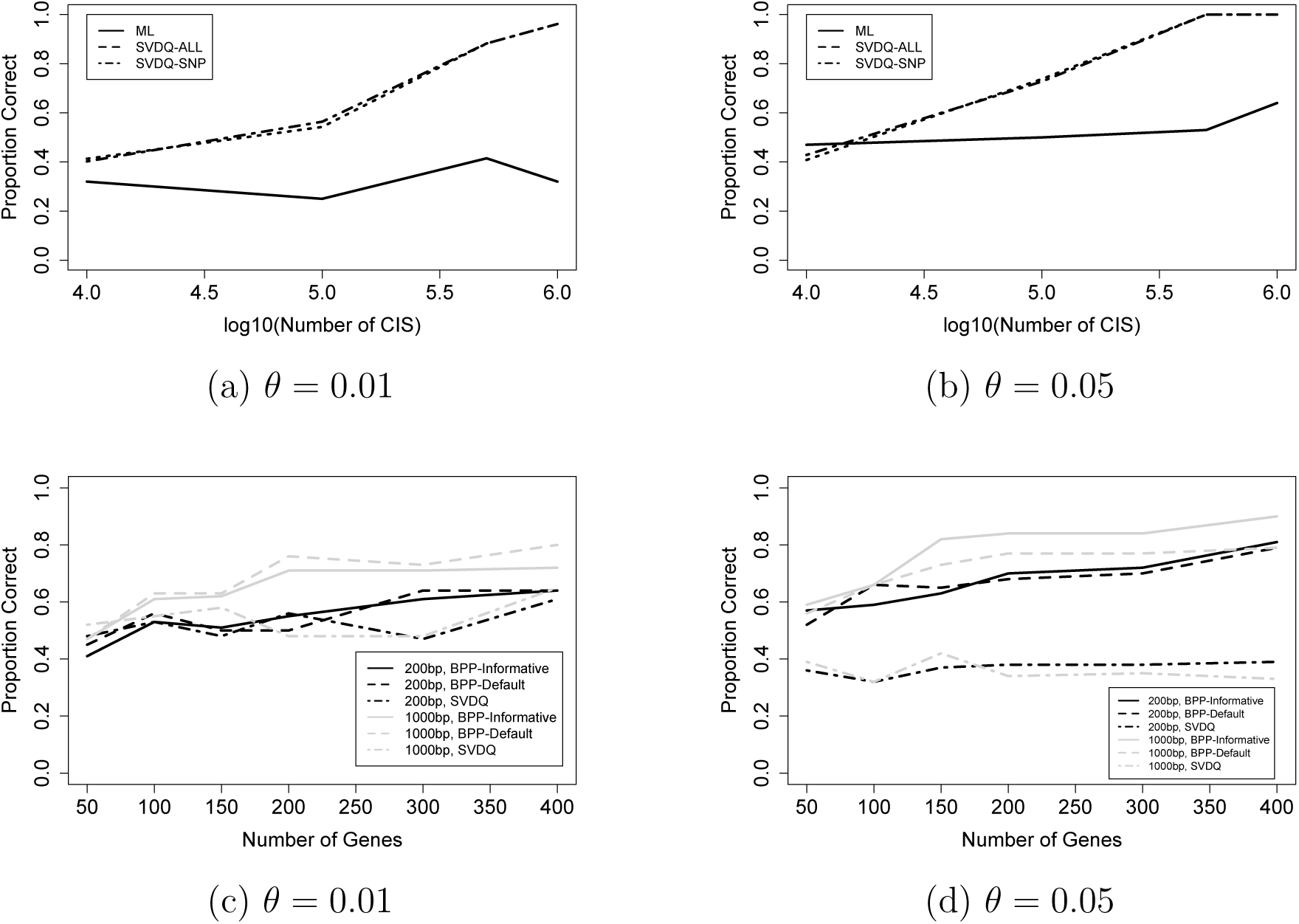
Results of the third simulation study which considers the anomalous species tree of Xu and Yang (2016) given by (Species1:0.48,(Species2:0.44, (Species3:0.4,Species4:0.4):0.04):0.04). In each plot, the x-axis shows the amount of data, and the y-axis shows the proportion of correctly-estimated unrooted species trees for each method.The first row shows the results for CIS data for (a) *θ* = 0.01 and (b) *θ* = 0.05. The second row shows the results for multilocus data for (c) *θ* = 0.01 and (d) *θ* = 0.05.

For the multilocus setting, we compared the performance of BPP with both informative and default priors, and we note that the choice of prior has an impact on the resulting inference. This is particularly apparent when *θ* = 0.05 (Figure 5(d)), where we see that the proportion correct decreases by ∼ 10% when default priors are used rather than priors centered at the true value of *θ*. This is because this particular choice of *θ* is larger than the typically-observed empirical values over which the default priors are centered. We selected this value in attempt to reproduce the high accuracy of BPP reported by Xu and Yang (2016) for this species tree. However, we also considered the more realistic value of *θ* = 0.01 (Figure 5(c)), where we see that the effect of the prior is less substantial, likely because the default prior puts more weight close to this value. However, BPP’s accuracy decreases for this value of *θ*, which can be attributed to the fact that the mutation rate is lower, resulting in fewer informative sites and therefore less information available for inference. Also notable is the effect of locus length in Figure 5(c) and (d), with shorter loci resulting in lower accuracy. Only with informative priors and the larger value of *θ* is accuracy substantially above 60% achieved for BPP when loci are 200bp in length. The strong performance of BPP noted by Xu and Yang (2016) for this tree is only achieved with a large value of *θ*, informative priors, and long loci (1000bp each).

We note that the performance of SVDQuartets is poor overall for the multilocus setting, with accuracy only around 50% for most conditions examined. Though this is similar to BPP’s accuracy for default priors, short loci, and a lower value of *θ*, all of which reflect common empirical conditions, it is clear that an increase in information available, either through longer loci or a larger value of *θ*, benefits BPP more than SVDQuartets. This result, together with the encouraging results for the CIS and SNP cases in Figure 5(a) and (b), further support our assertion that SVDQuartets may be most effective when the amount of data available, whether multilocus or SNP, is very large – precisely the situations in which Bayesian methods become more computationally expensive.

## SVDQuartets for Multilocus Data: 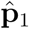 vs. 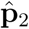

SVDQuartets was originally formulated for CIS data, and is easily applied to SNP data, as the constant patterns present in CIS data and absent in SNP data do not impact the reduced-rank results that form the theoretical basis of the method (see Chifman and Kubatko (2015) for details). However, in many cases, multilocus data have been sequenced and are already available. Recall that Theorem 0.5 showed that SVDQuartets is consistent when using either of

**Estimator 1:** 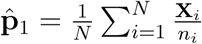

**Estimator 2:** 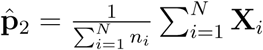
to estimate site pattern probabilities when multilocus data are generated via Definition 0.1. A natural question is then whether one of these estimators should be preferred. **?** examine this question and find that neither is uniformly better; rather the relative performance depends on the distribution *F* in Definition 0.1. Very generally, when *F* is concentrated around some value, 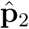 is better while when *F* is spread out, 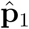 is better. For further discussion, we refer readers to **?**, noting that 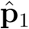 corresponds to their arithmetic average while 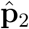 corresponds to their weighted average.

## Discussion

Our work gives the first consistency results for four-taxon species tree inference under the coalescent model for SVDQuartets for both CIS and multilocus data and for maximum likelihood for CIS data. Previous consistency results for maximum likelihood were only derived in the case of gene trees. In addition, we have proved that the SVDQuartets estimator has asymptotic error probability *O*(exp(−*CN*)) for CIS and multilocus data, where *N* is the number of loci. The constant *C* probably depends on the structure of the tree being estimated, but our simulations show that it does not appear to be particularly unreasonable in a variety of scenarios. We compare the performance of SVDQuartets and ML theoretically, and find that what is known rigorously is not sufficient to confirm the conjecture of Shi and Yang (2018) that ML is more efficient than SVDQuartets; rather the comparison is inconclusive, in large part because not enough is known about the performance of ML.We can only note that our error bounds for SVDQuartets and those conjectured from partial theoretical results for ML both take the form *O*(exp(−*CN*)) where the constant *C* may differ between the methods.

In our simulations, we assumed that the effective population size, *θ*, was constant throughout the tree. However, for empirical data, *θ* may vary from branch to branch, or even along branches within the tree. It is therefore important to note that our proofs of consistency did not rely on the assumption of constant effective population size. In the case of consistency of ML for CIS data, identifiability is known to hold when *θ* varies through the tree (Long and Kubatko, 2019) and expressions analogous to that in Equation (9) can be obtained for varying effective population sizes (Rusinko, 2018). In the case of the consistency of SVDQuartets, recent work (Long and Kubatko, 2019) has established that the method holds in the case of varying *θ*s, as well as in the absence of a molecular clock. Thus, the consistency result for SVDQuartets applies to a wide variety of mechanisms for data generation.

Our simulations demonstrate comparable performance for both ML and SVDQuartets for CIS data, while ML (as implemented in BPP) generally performs better with multilocus data. Importantly, our first simulation shows that SVDQuartets can be applied to SNP data without any loss of power to infer the true species tree, making it a good choice for computationally efficient analysis of SNP data under the MSC. Examination of the performance of these methods in the anomaly zone indicates that BPP can be sensitive to the choice of prior and to the number of sites within the loci, while SVDQuartets may require a large number of loci to obtain high accuracy. We also note that, at present, BPP implements only the JC69 model, while the theoretical results underlying SVDQuartets hold for the general time reversible (GTR) model and all submodels (Chifman and Kubatko, 2015), as well as for species trees that violate the molecular clock (Long and Kubatko, 2019), making the method quite generally applicable. Given the consistency results derived here, we suggest that for multilocus data, SVDQuartets will be a useful alternative to Bayesian methods such as BPP when the size of the data, in terms of the number of loci and/or the number of species, makes MCMC-based methods computationally prohibitive, i.e., our results indicate that SVDQuartets can be used to achieve consistent estimates of the species tree topology in precisely the cases in which Bayesian methods are currently computationally expensive.

## Acknowledgements

We thank Edward Susko, Ziheng Yang, and an anonymous reviewer for helpful comments on earlier drafts of this manuscript that led to its improvement. We are particularly grateful to Dr. Susko for suggesting a correction to our proof of consistency of SVDQuartets, and for several helpful comments about our overall approach.

